# A novel environmental DNA (eDNA) sampling method for aye-ayes from their feeding traces

**DOI:** 10.1101/272153

**Authors:** Megan L. Aylward, Alexis P. Sullivan, George H. Perry, Steig E. Johnson, Edward E. Louis

## Abstract

Non-invasive sampling is an important development in population genetic monitoring of wild animals. Particularly, the collection of environmental DNA (eDNA) which can be collected without needing to encounter the target animal, facilitates the genetic analysis of cryptic and threatened species. One method that has been applied to these types of sample is target capture and enrichment which overcomes the issue of high proportions of exogenous (non-host) DNA from these lower quality samples. We tested whether target capture of mitochondrial DNA from sampled feeding traces of wild aye-ayes would yield mitochondrial DNA sequences for population genetic monitoring. We sampled gnawed wood from feeding traces where aye-ayes excavate wood-boring insect larvae from trees. We designed RNA probes complementary to the aye-aye’s mitochondrial genome and used these to isolate aye-aye DNA from other non-target DNA in these samples. We successfully retrieved six near-complete mitochondrial genomes from two sites within the aye-aye’s geographic range that had not been sampled previously. This method can likely be applied to alternative foraged remains to sample species other than aye-ayes. Our method demonstrates the application to next-generation molecular techniques to species of conservation concern.

## Introduction

Genetic sampling of wild populations can help us to address questions of demography, individual relatedness, population structure, and other important aspects of biodiversity that cannot be answered by behavioural monitoring alone (Allendorf, Hohenlohe, & Luikart, 2010). However, the practicality of sampling wild animals can be limited by ethical implications due to the implicit risks of sedating animals to collect blood or tissue samples, especially for arboreal species, and for rare or elusive species (including nocturnal animals), by infrequent encounter rates. (Kohn & Wayne, 1997; Goossens, Chikhi, Utami, de Ruiter, & Bruford, 2000). Even when it is possible and safe to collect invasive samples from wild individuals, the number of samples that can be obtained with this approach may be fewer than analytically desirable (Sikes & Gannon, 2011). Under these conditions, non-invasive sampling is a valuable tool in advancing our understanding of wild populations.

Over the last two decades, advances in molecular biology have facilitated the use of non-invasive sampling for genetic analyses of wild populations (Morin, Wallis, Moore, Chakraborty, & Woodruff, 1993; Joost et al., 2007). Furthermore, traces of DNA left in the environment from biological material (eDNA) can be collected without even encountering target individuals. Source materials for non-invasive or eDNA sampling that have been used in wildlife monitoring and forensic studies include feathers, faeces, egg shells, hair, saliva, urine, and snail trails (Valiere & Taberlet, 2000; Inoue, Inoue-Murayama, Takenaka, & Nishida, 2007; Beja-Pereira, Oliveira, Alves, Schwartz, & Luikart, 2009; Sastre, Francino, Sanchez, & Ramirez, 2009). Non-invasive and eDNA sampling are particularly valuable in research surrounding unhabituated populations, cryptic species, and those at risk of extinction (Goossens et al., 2000; Cushman, McKelvey, Hayden, & Schwartz, 2006; KendallL et al., 2009; Clevenger & Sawaya, 2010; Schubert et al., 2011; Morin, Kelly, & Waits, 2016; Orkin, Yang, Yang, Yu, & Jiang, 2016).

One species that could especially benefit from the advances in non-invasive genetic sampling is the aye-aye (*Daubentonia madagascariensis*), which is a rare and elusive Malagasy primate of major conservation concern (Sterling & McCreless, 2007; Schwitzer et al., 2013). The low population density, large home range size, cryptic nature, and nocturnal activity of aye-ayes make them particularly difficult to locate, and precise distributions and population densities are unclear (Sterling, 1994b). Challenges of monitoring aye-ayes and obtaining reliable population dynamics data are reflected in the volatility of aye-aye’s conservation status designations over the last 70 years. In the 1950’s, the aye-aye was thought to be extinct (Sterling, 1994b). After aye-ayes were rediscovered in 1957, they were classified as Endangered; in 2008, their status was changed to Near Threatened before being reassessed as Endangered in 2012 (Andriaholinirina et al., 2015). In addition to being listed as Endangered, aye-ayes are currently considered one of the world’s top 25 most endangered primates (Randimbiharinirina et al., 2017 in Schwitzer et al. 2016-2018). Few encounters, along with the aye-aye’s solitary social organization and long maternal investment suggest low population densities. Low nuclear genomic diversity in aye-ayes reflects these assumptions; genomic analyses estimates of heterozygosity of 0.051%, and genetic diversity across synonymous sites of π= 0.073, are the lowest of any primate species studied to date (Perry, Melsted, et al., 2012; Perry, Reeves, et al., 2012). Therefore, despite the wide distribution of aye-ayes, there are likely few individuals, increasing the risk of local and global extinction (Schwitzer et al., 2013; Gross, 2017).

The IUCN’s lemur survival strategy recognises the need for biological monitoring of aye-ayes to better assess population status and conserve genetic diversity within this lineage (Schwitzer et al., 2013). In addition to their elusive behaviour making aye-ayes difficult to find and monitor, genetic sampling of individuals once they have been located is challenging; invasive collection of blood and tissue can only be achieved during immobilization, which is risky and must be conducted by trained and experienced personnel (Cunningham, Unwin, & Setchell, 2015). Therefore, to assess genetic diversity and identify priority populations for conservation a new, reliable means of non-invasive sampling in aye-ayes is required.

Reliable genotyping from non-invasively collected material holds much promise in sampling threatened species. Environmental DNA samples that are degraded due to exposure to both biotic and abiotic factors or contain high proportions of exogenous DNA can now provide reliable genetic markers (Beja-Pereira et al., 2009; Carpenter et al., 2013; Snyder-mackler et al., 2016). Techniques such as high-throughput sequencing technologies reduce sequencing errors and improve genotyping accuracy by providing greater depth of coverage across loci. One particularly promising approach is target capture, which provides a means of isolating endogenous DNA from the high proportions of exogenous DNA in non-invasive samples (Bi, Linderoth, Vanderpool, Good, & Nielsen, 2013; Hawkins et al., 2015; Kirillova et al., 2015; Kistler et al., 2015; Mohandesan et al., 2017). Specifically, RNA probes which are complementary to particular regions or markers in the genome of the target organism are specially designed and synthesized (Gnirke et al., 2009). These probes hybridize to the endogenous DNA in the sample. After hybridization, the biotin coating of the probes allows streptavidin-coated magnetic beads to bind; the bound probes and hybridized endogenous DNA can then be isolated from the exogenous DNA by using a magnet (Gnirke et al., 2009; Giolai et al., 2016). These developments make monitoring and sampling of wild populations where individuals are difficult to locate increasingly feasible.

For primates, the application of target capture from low quality samples has largely been applied to ancient DNA studies, but Perry et al., (2010) and Snyder-Mackler et al., (2016) have presented effective approaches for sampling endogenous DNA from faeces in extant primate species. Recently, Chiou & Bergey,(2018) demonstrated a different method that isolates endogenous DNA from primate faeces by targeting the higher CpG-methylation density of vertebrate taxa relative to the exogenous bacterial and plant DNA in faeces. Although faecal collection is a popular source of non-invasive sampling of primates (Oka & Takenaka, 2001; Quéméré, Crouau-Roy, Rabarivola, Louis, & Chikhi, 2010; Chancellor et al., 2011), it is unlikely to be a feasible method of sampling for aye-ayes. Factors such as aye-aye’s nocturnal behaviour, the height that they travel in the canopy, and their large nightly travel distances mean that defecation is difficult to observe and locating and collecting faecal material is problematic (Randimbiharinirina et al., 2017); accordingly, at two sites where aye-ayes have been monitored by Madagascar Biodiversity Partnership since 2010, aye-aye faecal samples have only been collected through routine immobilizations. Therefore, to gain information on the genetics of aye-aye populations to meet the IUCN aims, in this paper, we explore the possibility of sampling eDNA from aye-ayes (Schwitzer et al., 2013).

One potential source of eDNA in aye-ayes is from their distinct feeding traces left on trees. These traces are associated with their adaptations for extracting the larvae of wood-boring insects from tree trunks and branches, after identifying suitable foraging locations via a process of sniffing, lightly tapping, and listening (Sterling, 1994a; Erickson, 1995). Aye-ayes gnaw into selected areas of trees with their elongated and continuously growing incisors and extract larvae using their thin flexible third digit (Sterling, 1994a). During this foraging process, the buccal cavity of the aye-aye comes into contact with the wood (Sterling, 1994a; Erickson, 1995; Sterling & McCreless, 2007; arkive.org, 2017). Thus, trace amounts of aye-aye biological material in the form of epithelial cells of the buccal mucosa or from saliva may be deposited and accessible as an eDNA source.

We investigated the application of target capture and enrichment to obtain aye-aye DNA from aye-aye feeding traces. We aimed to determine whether this method is a feasible alternative to invasive sampling. If eDNA samples provide a means of remotely sampling wild aye-aye populations, we predict that (1) target enrichment is an effective method of obtaining aye-aye DNA from the exogenous DNA in feeding traces, and (2) we will be able to obtain full mitochondrial genomes (hereafter ‘mitogenomes’) for population genomic analysis.

## Methods

### Sampling sites and techniques

Initial training for researchers and local technicians on accurate identification of aye-aye feeding traces and the sample collection method was conducted through the Kianjavato Ahmanson Field Station (KAFS) located in the Kianjavato commune of the Vatovavy-Fitovinany region in southeast Madagascar during June 2014. KAFS is a long-term field site of the Madagascar Biodiversity Partnership (MBP). Individual aye-ayes present in the region have been fitted with ATS (Applied Telemetry Systems^®^) VHS radio-collars for behavioural monitoring, which afforded the opportunity to view known feeding sites in an area of bamboo to become familiar with density and frequency of feeding traces, their characteristics and to practice the collection technique (see below).

To then test the sampling method on unmonitored populations, we selected two study sites near to the limits of the aye-aye’s geographic distribution, where aye-ayes have been sighted but not sampled: Manombo Special Reserve in the Atsimo-Atsinanana region in southeastern Madagascar, operated by Madagascar National Parks and at that time supported by Durrell Wildlife Conservation Trust, and the Tsingy de Beanka, Ambinda in the Melaky region in west-central Madagascar and supported by Biodiversity Conservation Madagascar (Figure 1). Between July and December 2014, MA surveyed the forest at each site with the assistance of three local technicians. Prior to sampling, local technicians cleared undergrowth from existing trail systems in the forest to use as transects. During daylight hours, these transects were walked at a slow pace of approximately 1 km/hr to look for feeding traces and searched areas on either side of the transect at 200 m intervals. Traces were observed in both live and dead plants where characteristic holes gnawed by aye-aye were identified. These were confirmed by the distinct shape of feeding traces and typically presence of teeth marks (Figure 2). The research permit (N° 162/14/MEF/SG/DGF/DCB.SAP/SCB) for sample collection was obtained from the Ministere de l’environment, des eaux et forets et du tourisme, in Madagascar.

**Figure 1.**
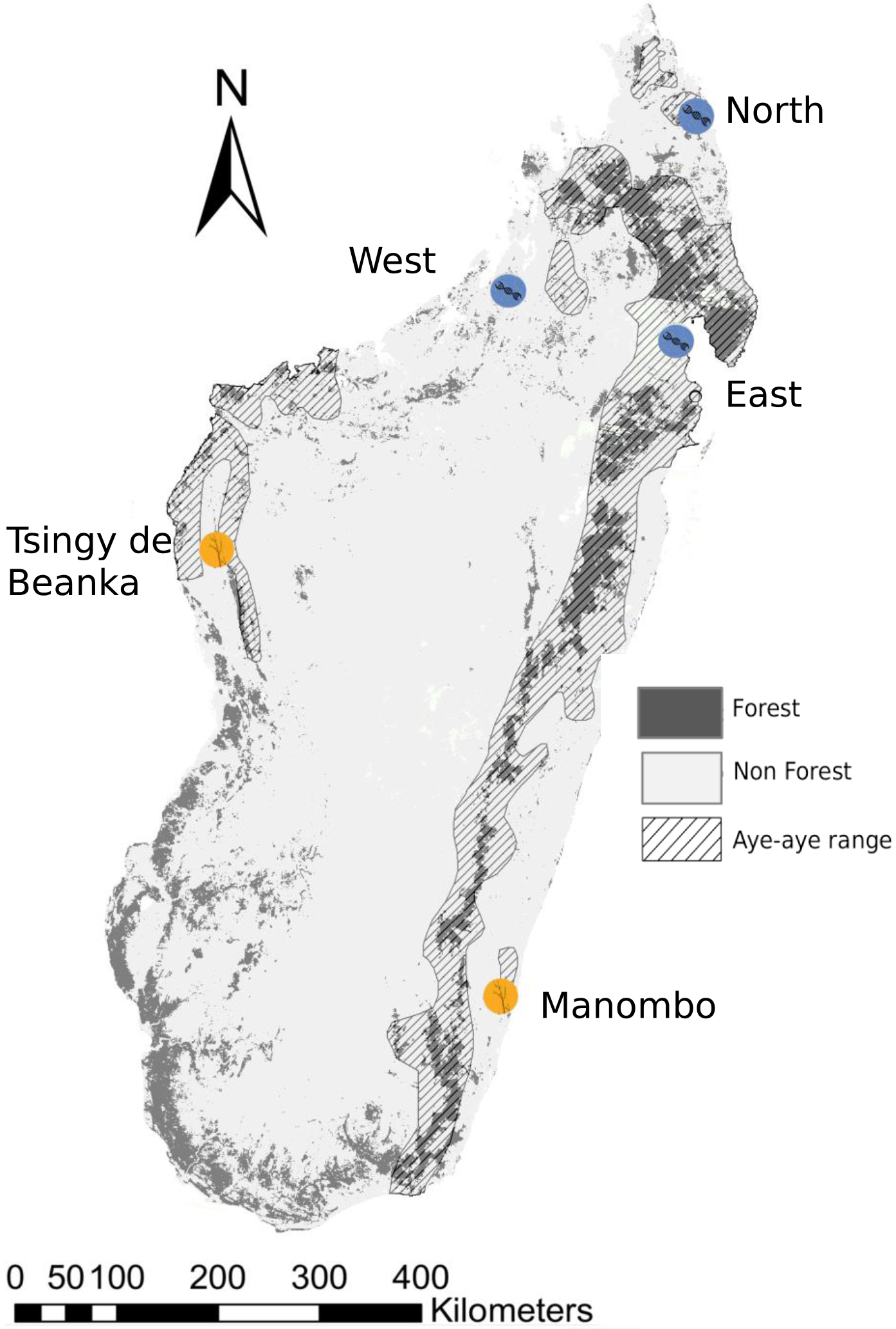
A map of the island of Madagascar showing sample locations targeted in this study. The orange circles highlight the two locations surveyed for aye-aye feeding traces and eDNA samples were collected. The blue circles indicate the locations that were sampled in Perry et al. 2013, from which the mitogenome sequences from Kistler et al. 2015 were sampled.

To collect samples, we selected areas around the edge of the trace where the buccal cavity and teeth of the animal were likely to have come into contact with the tree (Figure 2). MA took 3-6 samples of tree material per trace; each sample was approximately 2g^−1^ of wood shavings. MA collected samples using a scalpel to cut wood from the site directly into a 1.5 mL Eppendorf^®^ tube containing 500 μL RNAlater. To reduce the chance of contamination, MA wore latex gloves and used a new, sterile scalpel blade for each sample. At Manombo, we walked 9.5 km of transects over a 6 km2 area. At Beanka, we surveyed 10.8 km of transects covering a total 9 km^2^ area. MA sampled a total of 59 traces across the two sites (Manombo: n=27; Beanka: n=32). We stored samples at ambient temperature until they could be transferred to 4 °C for storage in Antananarivo, Madagascar. MBP exported samples from Madagascar (Export Permit Number 233N-EV07/MG16) and stored the samples at Omaha’s Henry Doorly Zoo and Aquarium at 4 °C for four months prior to processing.

**Figure 2.**
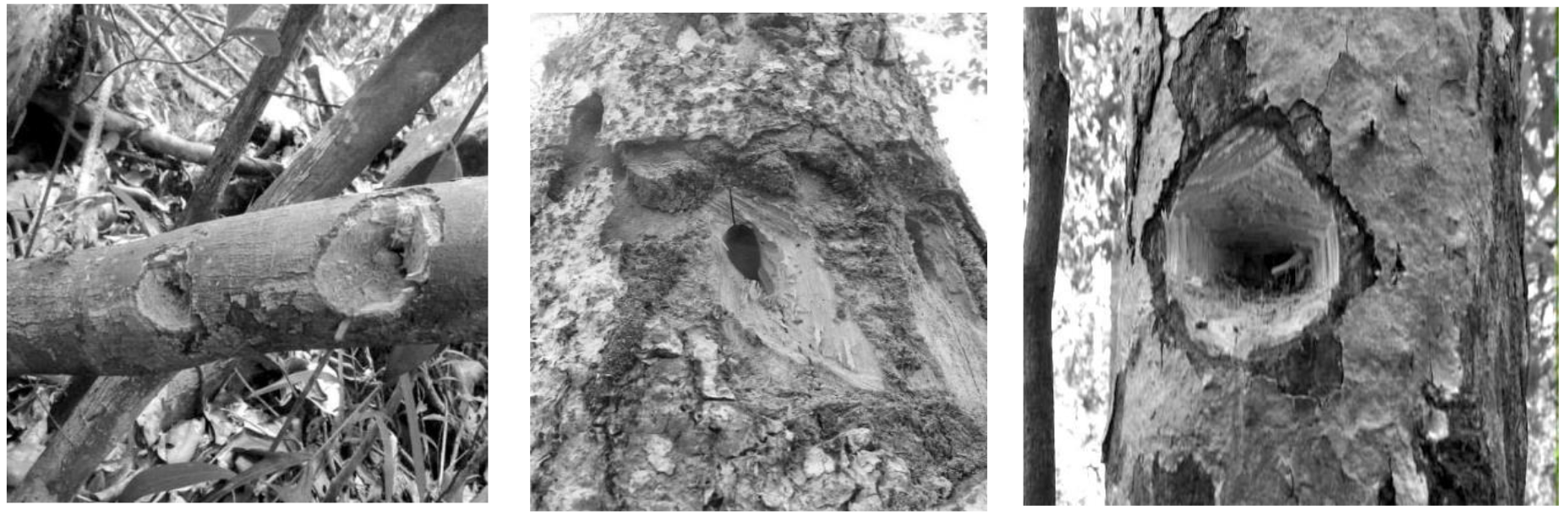
Aye-aye feeding traces. These images indicate the typical aye-aye feeding traces showing distinct shapes and teeth marks. These traces centre on a hole where the aye-aye has used its specialized extractive foraging strategy to access wood boring insect larvae. We sampled pieces of wood around the edge of the trace where the buccal cavity of the aye-aye comes into contact with the tree.

### DNA extraction, library preparation, capture, and sequencing

After samples were removed from storage, we extracted DNA from all samples. We vortexed each sample for 30 seconds in PBS to wash aye-aye DNA from the surface of the wood and then removed pieces of wood using sterile forceps. We added 500 μL of PBS solution to the sample and centrifuged at 10,000 g to pellet the cellular DNA and then carefully removed the supernatant so as not to disturb the precipitate. We performed all DNA extractions on the precipitate using EZNA^®^ blood DNA mini kit following the buccal swab protocol with the following modifications: we lysed samples overnight, and we eluted DNA in 50 μL elution buffer, incubated for 20 minutes and then re-eluted with the same eluate. Preliminary tests to confirm the presence of aye-aye DNA were conducted using a microsatellite marker for loci AYE33 of approximately 279 bp, with the following primer pair: forward 5’-3’ GTCTGCTACTCCTTAGGTGCTG and reverse 5’-TGGCTCAGGGCAATACAAT-3’. We amplified the extracted DNA in a 15 μL reaction containing 0.25 μL of each 10 uM primer, 7.5 μL buffer containing 1.5 mM MgCl_2_ (1X) and 0.2 mM of each dNTP (1X), 0.12 μL KAPA3G plant DNA polymerase, 4.9 μL H2O, and 2 μL DNA sample. Cycling parameters began with an initial enzyme activation temperature of 95 °C for 3 min, followed by 32 cycles of: denaturation at 98°C for 20 s, primer annealing at 58 °C for 15 s and elongation at 72 °C for 30 s, then one cycle of final elongation at 72 °C for 1 min. We visually confirmed the presence of the expected 279 bp PCR product by gel electrophoresis with 5 μL of PCR product and using 2 μL of a 1 kb ladder.

Extracted DNA was transported to Pennsylvania State University, University Park, PA for library preparation and DNA capture. We quantified the DNA concentration and fragment size of ten samples using a bioanalyzer (Agilent Technologies, Santa Clara, CA) at the Huck Institute of the Life Sciences (HILS), University Park, PA, genomics core facility. We selected a subset of 22 samples that were estimated to have been between 14 h-6 months old at the time of collection based on observations during surveys and discolouration of wood. We prepared DNA sequencing libraries from each of these samples using the Illumina TruSeq nano^®^ kit. The TruSeq nano^®^ library preparation protocol recommends 200 ng^−1^ of input DNA. Due to the typically low quantities of DNA extracted from each individual wood sample (<300 pg/uL), we combined DNA extracted from multiple samples taken from the same trace into one sample per feeding trace to increase total amounts of library preparation input DNA per sample to approximately 5-42 ng^−1^ DNA (based on bioanalyzer readings for a subset of samples). To account for the low input DNA quantities, we adapted the protocol to use half quantities of all reagents and volumes. We sheared each DNA sample to a mean size of 500 bp with the Covaris-M220^®^ using the following settings: 34 seconds, peak incident of 50, duty factor of 20% and 200 cycles per burst. For library preparation wash steps, 180 μL ethanol was used and an additional initial wash step was included to remove residual salts from eluting extracted DNA. We used the Bioanalyzer (Agilent Technologies, Santa Clara, CA) to quantify the final library and confirm a mean library size of 750 bp fragments, as expected for 550 bp libraries once the sequencing adapters and unique barcodes are attached to the 550bp DNA fragments.

As a preliminary test to check for the presence of aye-aye DNA in these libraries, we shotgun sequenced ten DNA libraries. Libraries were included as part of a larger multiplexed sequencing pool (Table A1) and sequenced using the Illumina NextSeq^®^ 500 2×150 pair end chemistry at UCLA Clinical Microarray Core Facility. Samples were demultiplexed at the sequencing facility.

RNA probes for capture of the aye-aye mitogenome were designed through MyBaits^®^ at Arbor Biosciences™. Baits were designed based on the aye-aye reference mitochondrial DNA genome sequence (GenBank^®^ accession NC_010299.1), bait length was 80 bp at 4x tiling and these are available as a predesigned MyBaits^®^ Mito panel. We conducted captures as per MyBaits^®^ manual v.3 and a single round of captures was conducted on 19 of the eDNA (foraging remains) samples. Post capture products were quantified using Bioanalyzer (Agilent Technologies, Santa Clara, CA) at the genomics core facility at the HILS and pooled for sequencing (Table A2). These capture products were sequenced as one pool of 17 samples and two of the samples (BTS38 and BTS108) were included as part of a different sequencing pool. The final mitochondrial capture pool of 17 Mito Bait captured DNA libraries was sequenced on one lane of the NextSeq 500 mid-output 2×150 bp pair end chemistry UCLA Clinical Microarray Core Facility. Samples were demultiplexed and barcodes removed at the sequencing facility.

## Data analysis

All computational analyses were conducted on servers provided by WestGrid (www.westgrid.ca) at Compute Canada (www.computecanada.ca). Raw reads were submitted to the NCBI SRA project under the accession PRJNA434884. The following processing and alignment pipeline was used for the shotgun and captured DNA sequencing pools. We assessed sequence read quality using Fast-QC (Andrews, 2007) and filtered reads based on minimum read length of 120bp and mean quality of 20 using PRINSEQ-lite (Schmieder & Edwards, 2011). We removed any potential human contaminant reads using the bbsplit package of bbmap (Bushnell, 2016); sequencing reads were aligned to both the aye-aye and human genome and any reads which matched better to the human genome than the aye-aye genome were removed from the dataset. Paired reads were aligned to the aye-aye reference genome GCA_000241425.1 (Perry, Reeves, et al., 2012), using the bwa-mem alignment algorithm (Li & Durbin, 2009; Li, 2013). We used the MarkDuplicates tool in PicardToolsv.1106 (Broad Institute, 2017) to remove sequencing and PCR duplicates, and reads flagged as clipped were filtered from the dataset. We generated consensus sequences for the eDNA samples with near-complete mitogenomes using samtoolsv.1.3, bcftoolsv.1.3 and vcfUtilsv.1.3 (Li et al., 2009; Danecek et al., 2011; Li, 2011). To confirm the base calls in these consensus sequences, we compared the samtoolsv.1.3 generated sequences to those generated using IGV (Robinson et al., 2012) to ensure the same nucleotide bases were found by each software program.

We compared the mitogenome sequences from our eDNA samples with the GenBank^®^ aye-aye mitogenome data to confirm that they were novel (Perry et al., 2013; Kistler et al., 2015). We called variants using GATK (McKenna et al., 2010), and used haplotype caller to generate individual gVCFs and then the joint variant caller. To filter for biallelic SNPs only, we used VCFtoolsv.1.12 and set maximum and minimum alleles to two. We used PGDSpiderv2.3 to convert SNP variant calls from vcf to ped format (Lischer & Excoffier, 2012). We used the --cluster and --matrix commands in Plinkv.1.7 (Purcell et al., 2007) to calculate the pairwise proportion (identity by state, IBS) of shared alleles at SNP loci among all complete mitogenome sequences. For the Plink analysis, we used the complete mitochondrial genomes excluding the hyper-variable region given the potential for homoplasy in this region and the possibility for multiple mutations per site (Ballard & Whitlock, 2004; Gonder, Mortensen, Reed, De Sousa, & Tishkoff, 2007).

## Results

### Pre-capture processing

After library preparation, shotgun sequencing of libraries from eDNA samples did not yield sufficient mitochondrial DNA for mitogenome analysis. Of the 10 libraries that were shotgun sequenced, seven yielded fewer than 10 reads aligning to the aye-aye mitochondrial genome; one sample, with the highest number of mitochondrial reads, was MSR58 with 189 unique reads (0.000274%) (Table 1).

**Table 1.** The fold increase in percent of unique on-target reads aligning to the aye-aye mitochondrial genome target MitoBait captures compared to shotgun sequencing of the DNA libraries. Data show for the ten libraries that were shotgun sequenced prior to capture.

| Sample ID | Number of on-target reads | Unique on-target reads, shotgun sequencing (%) | Unique on-target reads, from captured DNA libraries (%) | Fold enrichment |
| --- | --- | --- | --- | --- |
| MSR01 | 0 | 0.000000 | 0.0027 | - |
| MSR44 | 8 | 0.000038 | 0.0538 | 1416 |
| MSR46 | 0 | 0.000000 | 0.0019 | - |
| MSR50 | 31 | 0.000063 | 0.0248 | 394 |
| MSR58 | 189 | 0.000274 | 0.0408 | 149 |
| BTS04 | 0 | 0.000000 | 0.0051 | - |
| BTS38 | 6 | 0.000103 | 0.0267 | 259 |
| BTS60 | 0 | 0.000000 | 0.0005 | - |
| BTS108 | 24 | 0.000124 | 0.0139 | 112 |
| BTS112 | 0 | 0.000000 | 0.0056 | - |

### Target capture DNA sequencing

Target captures of the mitogenome increased the proportion of on-target mitochondrial DNA reads by several orders of magnitude compared to reads yielded from shotgun sequencing without capture (Table 1). For samples for which it was possible to calculate the fold enrichment (i.e., the samples that were both shotgun sequenced and sequenced following DNA capture), the percent of unique on-target reads increased by over 100 times. For captured samples, the percent of on-target reads ranged from 0.009% to 27.5% of the total reads, whereas the percent unique on-target reads ranged from 0.001% to 0.05% (Table 2).

**Table 2.** Sequencing summary statistics showing number of reads and depth and breadth of coverages for MitoBait captures. Grey shading indicates samples with over 90% breadth of coverage that were used for comparisons with published mitogenomes.

| Sample site | Sample ID | Reads on-target (%) | Unique reads on-target (%) | Mean read depth (SD) | Mean % coverage |
| --- | --- | --- | --- | --- | --- |
| Tsingy de Beanka | BTS04 | 0.06 | 0.0015 | 1.56 (0.9) | 59.3 |
| Tsingy de Beanka | BTS27 | 0.38 | 0.0063 | 13.0 (5.3) | 99.4 |
| Tsingy de Beanka | BTS38 | 0.22 | 0.027 | 23.3 (5.9) | 98.0 |
| Tsingy de Beanka | BTS60 | 0.02 | 0.0005 | 1.8 (2.2) | 45.6 |
| Tsingy de Beanka | BTS68 | 4.3 | 0.0088 | 8.5 (6.0) | 98.9 |
| Tsingy de Beanka | BTS102 | 0.26 | 0.0013 | 1.8 (1.4) | 27.3 |
| Tsingy de Beanka | BTS105 | 0.5 | 0.0005 | 1.7 (1.2) | 22.1 |
| Tsingy de Beanka | BTS108 | 0.4 | 0.014 | 15.1 (5.6) | 99.5 |
| Tsingy de Beanka | BTS112 | 0.46 | 0.0056 | 1.3 (0.6) | 10.9 |
| Manombo Special | MSR01 | 0.05 | 0.0027 | 2.0 (1.4) | 65.9 |
| Manombo Special | MSR23 | 0.04 | 0.039 | 1.2 (0.4) | 12.3 |
| Manombo Special | MSR37 | 0.16 | 0.0002 | 1.1 (0.3) | 11.1 |
| Manombo Special | MSR39 | 27.54 | 0.0024 | 5.0 (11.83) | 45.5 |
| Manombo Special | MSR44 | 22.25 | 0.05 | 1.5 (1.2) | 21.9 |
| Manombo Special | MSR46 | 4.11 | 0.0019 | 1.3 (0.5) | 10.8 |
| Manombo Special | MSR50 | 12.19 | 0.025 | 2.5 (2.07) | 70.9 |
| Manombo Special | MSR56 | 0.024 | 0.0001 | 1.3 (0.6) | 15.0 |
| Manombo Special | MSR58 | 2.71 | 0.04 | 12.4 (7.2) | 98.5 |
| Manombo Special | MSR62 | 2.20 | 0.015 | 25.1 (8.8) | 99.8 |

We recovered near-complete mitogenomes for six of the 19 captured samples (31.6%). The depth and breadth of coverage varied between samples (Table 1). For these six mitogenomes, the mean depth of coverage was 16x (range 8-25x). For the 13 samples with less than 90% coverage, the percent of the genome covered ranged from 10-71% (mean 32.58%) with a mean depth of 1.7 (+/−0.3-2.8). Five of these 13 samples had a breadth of coverage between 45-70% and were all identical at SNP loci to the samples with greater breadth of coverage. We only included samples with coverage greater than 90% in downstream analysis.

### Mitogenome analysis

Based on multiple sequence alignment of these seven near-complete mitogenomes, we identified one unique mitogenome sequence from Manombo Special Reserve and two unique mitogenomes from Tsingy de Beanka. Alignment with previously published GenBank^®^ sequences that the mitogenome sequences obtained from the eDNA samples were unique to all previously published aye-aye mitogenomes. For the GenBank sequences sample locations from northern Madagascar, we follow the population names in Perry et al., (2013); North, East, and West (Figure 1). When we compared SNP loci among samples, all mitogenomes clustered into distinct populations according to their collection location (Figure 3). Pairwise comparison of genotypes at polymorphic loci indicate that there are four distinct populations (Figure 3). The Tsingy de Beanka population showed moderate differentiation from the other populations (Figure 3). There was also a split between the East population and Manombo in the southeast, while the North population shared the fewest alleles with all other populations.

**Figure 3.**
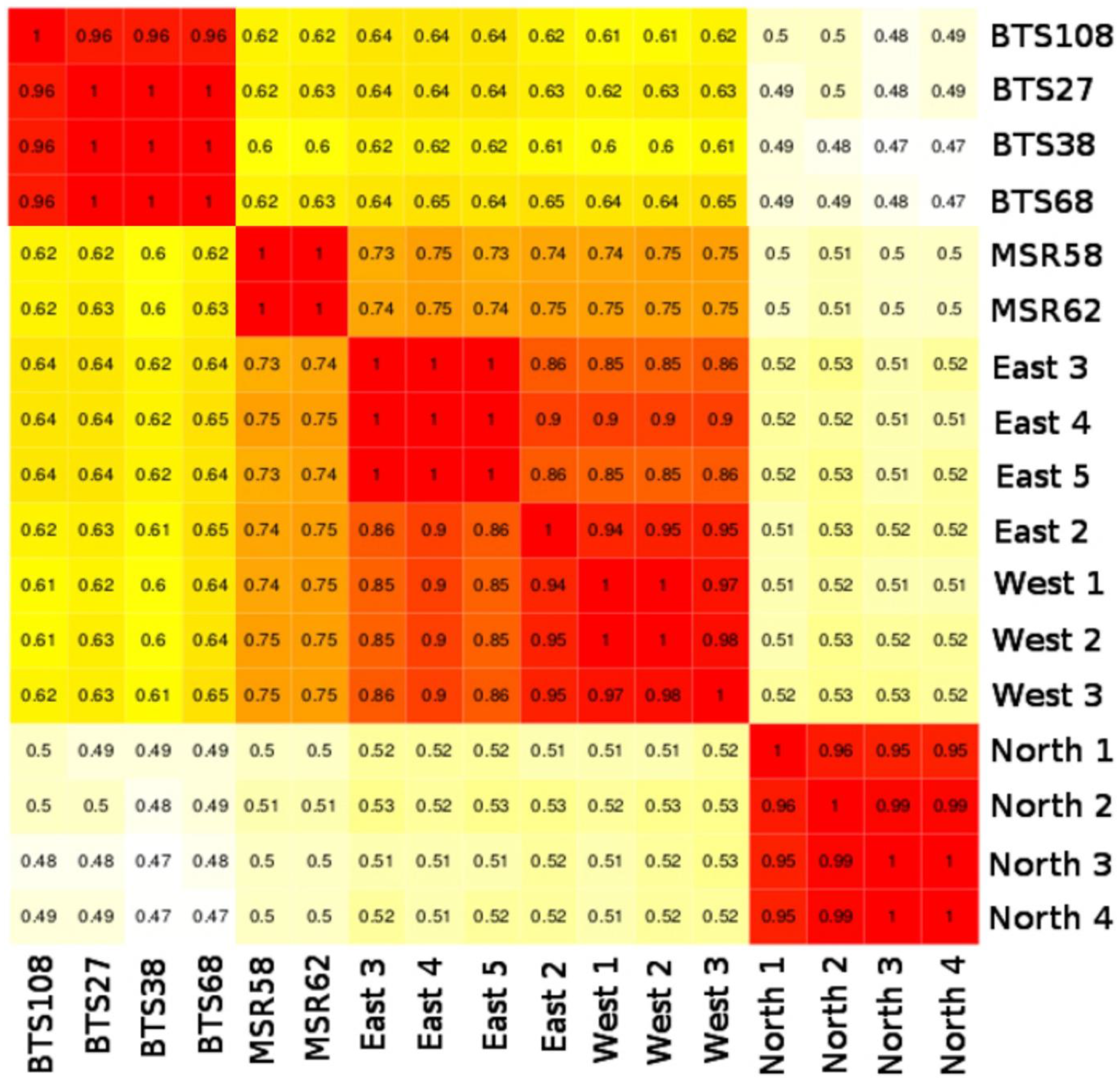
Identity by state matrix on the left, which shows the proportion of pairwise shared alleles at each SNP site for mitogenomes excluding the d-loop region. We compare eDNA samples from feeding traces with previously published mitogenome sequences (Kistler et al., 2015). Warmer colours indicate greater proportions of shared alleles at these loci.

## Discussion

We tested whether sampling of aye-aye feeding traces is a feasible method of non-invasive sampling of this species. Our first prediction, that target capture would provide a means of sampling aye-aye DNA from these feeding traces, was supported by the increased amount of endogenous DNA obtained from target capture compared to shotgun sequencing. The number of on-target reads from the shotgun sequencing was not at sufficient coverage across the mitogenome to identify polymorphic sites and genotype individuals. The high level of enrichment from captured libraries compared to shotgun sequencing indicates the efficacy of a target capture approach (Gnirke et al., 2009). Our second prediction, that mitogenomes could be obtained from these samples and used for population genomic analysis, was also supported. We were able to obtain near complete mitogenomes for 31.6% of the processed samples. Previous studies that have sampled eDNA from foraged material have typically only retrieved partial D-loop fragments rather than whole mitogenomes (Nichols, Konigsson, Danell, & Spong, 2012; Caniglia, Fabbri, Mastrogiuseppe, & Randi, 2013; Wheat, Allen, Miller, Wilmers, & Levi, 2016). While partial fragments can be useful in metabarcoding studies for species detection, whole mitogenomes provide greater phylogenetic resolution and reduce the likelihood of including nuclear copies of mitochondrial DNA(numts) (Johnsen, Kearns, Omland, & Anmarkrud, 2017).

### Future directions for methodological improvements for sampling aye-aye eDNA

The method developed here is a valuable tool to extend sampling of unmonitored aye-aye populations. Yet, improved efficacy of the method to increase amount of aye-aye DNA obtained and therefore reduce the cost per sample could make it a more attractive application for conservation monitoring (Ekblom & Galindo, 2010). Low proportions of on-target, unique sequence reads recovered are common when working with environmental samples (Ávila-Arcos et al., 2015; Nielsen et al., 2017) and adequate sampling effort is required to account for this issue. A mode of three different samples was taken per feeding trace, and it would be feasible to double this number. Therefore, we recommend more intensive sampling of each feeding trace to allow for multiple library preparations per trace, whilst maintaining sufficient amounts of input DNA for TruSeq nano^®^ library preparation. Multiple library preparations for each trace prior to capture may increase the number of unique DNA fragments within the samples and reduce PCR bias. The proportion of unique reads aligning to the target region was relatively low (0.4-12 %). To better assess the number of PCR cycles to run post-capture to achieve sufficient concentrations for sequencing, qPCR quantification may be valuable to ensure reduced PCR duplication, therefore improved sequencing efficacy and ultimately increased cost effectiveness (Enk, Rouillard, & Poinar, 2013).

### Application towards addressing the IUCN’s objectives for aye-ayes

We demonstrated a novel method that can be applied across the species range for population genomic monitoring. This method helps to meet objective seven of the IUCN’s lemur survival plan set out by Schwitzer et al., (2013:page32) which is “Fill knowledge gaps in population ecology and biodiversity of lemurs, and increase training of Malagasy scientists”. Our method provides a means of genomic biodiversity monitoring using cutting edge molecular techniques. Application of this method allowed us to successfully recover six aye-aye mitogenomes from sites in the south-east and west of Madagascar where aye-ayes have not been sampled previously. The wide distribution of aye-ayes across Madagascar means sampling across environments is key in effective population monitoring at local and national scales (Schwitzer et al., 2013). The different habitat types that aye-ayes are distributed in range from rainforests in southeast Madagascar, a habitat with relatively humidity and rainfall, to the dry deciduous forest in the west of Madagascar, with less dense canopy cover and likely greater exposure to direct UV radiation (Rakoto-Joseph, Garde, David, Adelard, & Randriamanantany, 2009; Andriamisedra, Aylward, Johnson, Louis, & Raharivololona, 2015). Our recovery of mitogenomes from distinct environments indicates these abiotic factors do not necessarily preclude the ability to sample eDNA from these areas and this method can likely be applied to sample in habitat across the species’ geographic distribution (Sterling & McCreless, 2007). The mobility of this sampling technique along with little training required for sample collection provides a means of surveying large areas of aye-aye habitat in a relatively short time frame. The ease of sampling and ability to survey large areas provide the opportunity for sampling multiple individuals across a population and accessing mitogenome diversity within forest fragments.

Our preliminary analysis revealed high differentiation between the North population and all other sampled locations, consistent with the whole genome analysis of Perry et al., (2013) which showed the North population to be the most genetically distinct of three regional populations. Given that our relatively sparse sampling strategy revealed two genetically distinct populations, we encourage the application of this method and the use of the predesigned aye-aye Mito panel from MyBaits^®^ to sample additional areas of aye-aye habitat.

### Applications to other species

The aye-aye is arguably one of the most elusive lemur species in Madagascar (Schwitzer et al., 2013), and other endangered species that are difficult to locate could also benefit from eDNA sampling. The Critically Endangered *Prolemur simus* leave traces on bamboo plants which are similar to those of sympatric bamboo lemurs that feed on the same plant species (Tan, 1999; Ravaloharimanitra et al., 2011). Application of the method presented here could confirm the presence of these threatened species which are typically sparsely distributed and difficult to locate in the wild (Wright et al., 2008; Ravaloharimanitra et al., 2011; Frasier et al., 2015). Similarly, Critically Endangered *Eulemur cinereiceps* could be sampled via traces on gnawed fronds of *Cecropia peltata* trees (Ralainasolo, Ratsimbazafy, & Stevens, 2006). Although the method developed here is novel, sampling of saliva from foraged material has been used previously as a means of non-invasive sampling; gorillas and golden monkeys discard plant material (Smiley et al., 2010), ungulates leave saliva traces on foraged twigs (Nichols et al., 2012), and large carnivores leave saliva on prey (Blejwas, Williams, Shin, Dale, & Jaeger, 2006; Sundqvist, Ellegren, & Vila, 2008; Glen et al., 2010; Caniglia et al., 2013; Wheat et al., 2016). This method of target capture could be applied to these sources to sample mitogenome eDNA from a range of species.

### NGS molecular techniques and conservation

This method provides an example of the utility of next-generation molecular techniques for non-invasive samples towards sampling wild populations in a species that requires conservation attention. The application of target capture and enrichment has been used previously in non-human primates to sample faecal and ancient DNA (Perry et al., 2010; Kistler et al., 2015; Snyder-mackler et al., 2016). We demonstrated the application of similar molecular techniques to sample from an Endangered lemur species, revealing broad-scale population structure across the aye-aye’s geographic range. One area of discussion that arises from reviews of the application of next-generation molecular techniques to conservation is often the necessity of these novel techniques over more conventional approaches, given the increased cost and amount of data generated (Allendorf et al., 2010; Mcmahon, Teeling, & Höglund, 2014; Shafer et al., 2015). Here we present both a unique challenge and solution to sampling a low-density, elusive, and Endangered species. These methodological developments are valuable tools that have enabled us to sample and monitor a cryptic species that otherwise has limited genomic sampling potential. As the field of conservation genomics expands methods such as the one presented here can be applied to achieve direct conservation action.

## Author contributions

EL, GP, MA and SJ conceived idea and designed methodology; MA collected samples; MA and AS conducted laboratory work. All authors contributed critically to drafts and gave final approval for publication.

## Data accessibility

Raw sequence reads are available from GenBank SRA accession SRP133213

## Supporting information

**Table A1.** Sequencing pool parameters for shotgun sequencing of eDNA libraries.

| Sample | Conc. ng/ul | Vol. in pool | Total ng | % pool by amount |
| --- | --- | --- | --- | --- |
| MSR01 | 0.26 | 2 | 0.528 | 6.81 |
| MSR44 | 0.17 | 5.88 | 0.999 | 6.58 |
| MSR46 | 0.46 | 2.17 | 0.999 | 6.58 |
| MSR50 | 0.39 | 2.56 | 0.998 | 6.58 |
| MSR58 | 0.96 | 1.04 | 0.994 | 6.58 |
| BTS04 | 0.53 | 1.89 | 1 | 6.6 |
| BTS38 | 0.03 | 1 | 0.03 | 0.2 |
| BTS60 | 0.70 | 1.43 | 1 | 6.59 |
| BTS108 | 0.20 | 1 | 0.2 | 1.32 |
| BTS112 | 0.34 | 2.94 | 0.999 | 6.58 |

**Table A2.** Sequencing pool parameters for MitoBait captures from eDNA samples.

| Sample | Conc. ng/ul | Vol. in pool | ng total | % pool by amount |
| --- | --- | --- | --- | --- |
| BTS04 | 0.36 | 11 | 3.96 | 6.7 |
| BTS27 | 6.8 | 3 | 4.08 | 6.9 |
| BTS60 | 0.515 | 7.8 | 4.02 | 6.8 |
| BTS68 | 0.633 | 6.3 | 3.99 | 6.75 |
| BTS102 | 4.4 | 0.8 | 3.52 | 5.96 |
| BTS105 | 1.56 | 2.6 | 4.06 | 6.87 |
| BTS112 | 0.23 | 16.7 | 2.87 | 4.86 |
| MSR01 | 0.43 | 9.4 | 4 | 6.77 |
| MSR23 | 0.546 | 7.3 | 3.99 | 6.75 |
| MSR37 | 0.389 | 10.3 | 4.01 | 6.79 |
| MSR39 | 0.848 | 4.7 | 3.9 | 6.75 |
| MSR44 | 0.003 | 10 | 0.03 | 0.05 |
| MSR46 | 0.003 | 10 | 0.03 | 0.05 |
| MSR50 | 0.0 | 10 | 0 | 0.00 |
| MSR56 | 0.932 | 4.3 | 4.01 | 6.79 |
| MSR58 | 0.209 | 10 | 2.09 | 3.54 |
| MSR62 | 2.66 | 1.5 | 3.99 | 6.75 |
| BTS108 | 0.42 | 6 | 2.5 | N/A |
| BTS38 | 2.2 | 1.8 | 3.96 | N/A |

## References

Allendorf, F. W., Hohenlohe, P. A., & Luikart, G. (2010). Genomics and the future of conservation genetics. Nature Reviews Genetics, 11(10), 697–709. doi:10.1038/nrg2844

Andriaholinirina, N., Baden, A., Blanco, M., Chikhi, L., Cooke, A., Davies, N.,…Zaramody. (2015). Daubentonia madagascariensis,. The IUCN Red List of Threatened Species, 8235. doi:10.2305/IUCN.UK.2014-

Andriamisedra, R., Aylward, M., Johnson, S. E., Louis, E. E., & Raharivololona, B. M. (2015). Détermination de quelques aspects de l’écologie de Daubentonia madagascariensis (Gmelin, 1788) dans deux forêts…Lemur News, 19. arkive.org. (2017). Aye-aye feeding on grubs using elongated finger. BBC Motion Gallery.

Ávila-Arcos, M. C., Sandoval-Velasco, M., Schroeder, H., Carpenter, M. L., Malaspinas, A. S., Wales, N.,…Gilbert, M. T. P. (2015). Comparative performance of two whole-genome capture methodologies on ancient DNA Illumina libraries. Methods in Ecology and Evolution, 6(6), 725–734. doi:10.1111/2041-210X.12353

Ballard, J. W. O., & Whitlock, M. C. (2004). The incomplete natural history of mitochondria. Molecular Ecology, 13(4), 729–744. doi:10.1046/j.1365- 294X.2003.02063.x

Beja-Pereira, A., Oliveira, R., Alves, P. C., Schwartz, M. K., & Luikart, G. (2009). Advancing ecological understandings through technological transformations in noninvasive genetics. Molecular Ecology Resources, 9(5), 1279–301. doi:10.1111/j.1755-0998.2009.02699.x

Bi, K. E., Linderoth, T., Vanderpool, D. A. N., Good, J. M., & Nielsen, R. (2013). Unlocking the vault: next-generation museum population genomics. Molecular Ecology, 22, 6018–6032. doi:10.1111/mec.12516

Blejwas, K. M., Williams, C. L., Shin, G. T., Dale, R., & Jaeger, M. M. (2006). Salivary DNA Evidence Convicts Breeding Male Coyotes of Killing Sheep. Journal of Wildlife Management, 70(4), 1087–1093.

Bushnell, B. (2016). BBMap short read aligner. University of California, Berkeley, California. URL Http://Sourceforge.Net/Projects/Bbmap.

Caniglia, R., Fabbri, E., Mastrogiuseppe, L., & Randi, E. (2013). Who is who? Identification of livestock predators using forensic genetic approaches. Forensic Science International: Genetics, 7, 397–404.

Carpenter, M. L., Buenrostro, J. D., Valdiosera, C., Schroeder, H., Allentoft, M. E., Sikora, M.,…Gilbert, M. T. P. (2013). Pulling out the 1 %: Whole-Genome Capture for the Targeted Enrichment of Ancient DNA Sequencing Libraries. The American Journal of Human Genetics, 93, 852–864. doi:10.1016/j.ajhg.2013.10.002

Chancellor, R. L., Satkoski, J., George, D., Olupot, W., Lichti, N., Smith, D. G., & Waser, P. M. (2011). Do Dispersing Monkeys Follow Kin? Evidence from Gray-cheeked Mangabeys (Lophocebus albigena). International Journal of Primatology, 32(2), 474–490. doi:10.1007/s10764-010-9483-6

Chiou, K. L., & Bergey, C. M. (2018). Methylation-based enrichment facilitates low-cost, noninvasive genomic scale sequencing of populations from feces. Scientific Reports, 8(1). doi:10.1038/s41598-018-20427-9

Clevenger, A. P., & Sawaya, M. A. (2010). Piloting a Non-Invasive Genetic Sampling Method for Evaluating Population-Level Benefits of Wildlife Crossing Structures. Ecology and Society, 15(1).

Cunningham, E. P., Unwin, S., & Setchell, J. M. (2015). Darting Primates in the Field: A Review of Reporting Trends and a Survey of Practices and Their Effect on the Primates Involved. International Journal of Primatology, 36(5), 911–932. doi:10.1007/s10764-015-9862-0

Cushman, S. a, McKelvey, K. S., Hayden, J., & Schwartz, M. K. (2006). Gene flow in complex landscapes: testing multiple hypotheses with causal modeling. The American Naturalist, 168(4), 486–99. doi:10.1086/506976

Danecek, P., Auton, A., Abecasis, G., Albers, C. A., Banks, E., Depristo, M. A.,…Vcf, T. (2011). The variant call format and VCFtools. Bioinformatics (Oxford, England), 27(15), 2156–2158. doi:10.1093/bioinformatics/btr330

Ekblom, R., & Galindo, J. (2010). Applications of next generation sequencing in molecular ecology of non-model organisms. Heredity, 107(1), 1–15. doi:10.1038/hdy.2010.152

Enk, J., Rouillard, J., & Poinar, H. (2013). Quantitative PCR as a predictor of aligned ancient DNA. Biotechniques. doi:10.2144/000114114

Erickson, C. J. (1995). Feeding Sites for Extractive Foraging by the Aye-Aye, Daubentonia madagascariensis, 240, 235–240.

Frasier, C. L., Rakotonirina, J.-N., Razanajatovo, L. G., Nasolonjanahary, T. S., Rasolonileniraka, Mamiaritiana, S. B.,…Louis, E. E. (2015). Expanding Knowledge on Life History Traits and Infant Development in the Greater Bamboo Lemur (Prolemur simus): Contributions from Kianjavato, Madagascar. Primate Conservation, 29(1), 75–86. doi:10.1896/052.029.0110

Giolai, M., Paajanen, P., Verweij, W., Percival-Alwyn, L., Baker, D., Witek, K.,…Clark, M. D. (2016). Targeted capture and sequencing of gene-sized DNA molecules. BioTechniques, 61(6), 315–322. doi:10.2144/000114484

Glen, A. S., Berry, O., Sutherland, D., Garretson, S., Robinson, T., & de Tores, P. J. (2010). Forensic DNA confirms intraguild killing of a chuditch (Dasyurus geoffroii) by a feral cat (Felis catus). Conservation Genetics, 11(September 2014), 1099–1101. doi:10.1007/s10592-009-9888-y

Gnirke, A., Melnikov, A., Maguire, J., Rogov, P., LeProust, E. M., Brockman, W.,…Nusbaum, C. (2009). Solution hybrid selection with ultra-long oligonucleotides for massively parallel targeted sequencing. Nature Biotechnology, 27(2), 182–189. doi:10.1038/nbt.1523

Gonder, M. K., Mortensen, H. M., Reed, F. A., De Sousa, A., & Tishkoff, S. A. (2007). Whole-mtDNA genome sequence analysis of ancient african lineages. Molecular Biology and Evolution, 24(3), 757–768. doi:10.1093/molbev/msl209

Goossens, B., Chikhi, L., Utami, S. S., de Ruiter, J., & Bruford, M. W. (2000). A multi-samples, multi-extracts approach for microsatellite analysis of faecal samples in an arboreal ape. Conservation Genetics, 1(2), 157–162. doi:10.1023/A:1026535006318

Gross, M. (2017). Primates in peril. Current Biology, 27(12), R573–R576. doi:10.1016/j.cub.2017.06.002

Hawkins, M. T. R., Hofman, C. A., Callicrate, T., Mcdonough, M. M., Tsuchiya, M. T. N., Ecer, E. L. I., & Errez, E. G. (2015). In-solution hybridization for mammalian mitogenome enrichment: pros, cons and challenges associated with multiplexing degraded DNA. Molecular Ecology Resources. doi:10.1111/1755-0998.12448

Inoue, E., Inoue-Murayama, M., Takenaka, O., & Nishida, T. (2007). Wild chimpanzee infant urine and saliva sampled noninvasively usable for DNA analyses. Primates, (48), 156–159. doi:10.1007/s10329-006-0017-y

Johnsen, A., Kearns, A. M., Omland, K. E., & Anmarkrud, J. A. (2017). Sequencing of the complete mitochondrial genome of the common raven Corvus corax (Aves: Corvidae) confirms mitogenome-wide deep lineages and a paraphyletic relationship with the Chihuahuan raven C. cryptoleucus. PLOS ONE, 12(10), e0187316. doi:10.1371/journal.pone.0187316

Joost, S., Bonin, a, Bruford, M. W., Després, L., Conord, C., Erhardt, G., & Taberlet, P. (2007). A spatial analysis method (SAM)to detect candidate loci for selection: towards a landscape genomics approach to adaptation. Molecular Ecology, 16(18), 3955–69. doi:10.1111/j.1365-294X.2007.03442.x

KendallL, K. C., Stetz, J. B., Boulanger, J., MacLeod, A. M. Y. C., Paetkau, D., & White, G. C. (2009). Demography and Genetic Structure of a Recovering Grizzly Bear Population. The Journal of Wildlife Management, 73(1), 3–16. doi:10.2193/2008-330

Kirillova, I. V., Zanina, O. G., Chernova, O. F., Lapteva, E. G., Trofimova, S. S., Lebedev, V. S.,…Shapiro, B. (2015). An ancient bison from the mouth of the Rauchua River (Chukotka, Russia). Quaternary Research, 84(2), 232–245. doi:10.1016/j.yqres.2015.06.003

Kistler, L., Ratan, A., Godfrey, L. R., Crowley, B. E., Hughes, C. E., Lei, R.,…Perry, G. H. (2015). Comparative and population mitogenomic analyses of Madagascar’s extinct, giant ‘subfossil’ lemurs. Journal of Human Evolution, 79, 45–54. doi:10.1016/j.jhevol.2014.06.016

Kohn, M. H., & Wayne, R. K. (1997). Facts from faeces revisited. TREE, 12, 223–227.

Li, H. (2011). A statistical framework for SNP calling, mutation discovery, association mapping and population genetical parameter estimation from sequencing data. Bioinformatics, 27(21), 2987–2993. doi:10.1093/bioinformatics/btr509

Li, H. (2013). Aligning sequence reads, clone sequences and assembly contigs with BWA-MEM.

Li, H., & Durbin, R. (2009). Fast and accurate short read alignment with Burrows-Wheeler transform. Bioinformatics, 25(14), 1754–1760. doi:10.1093/bioinformatics/btp324

Li, H., Handsaker, B., Wysoker, A., Fennell, T., Ruan, J., Homer, N.,…Durbin, R. (2009). The Sequence Alignment/Map format and SAMtools. Bioinformatics, 25(16), 2078–2079. doi:10.1093/bioinformatics/btp352

Lischer, H. E. L., & Excoffier, L. (2012). PGDSpider: an automated data conversion tool for connecting population genetics and genomics programs. Bioinformatics, 28(2), 298–298. doi:10.1093/bioinformatics/btr642

McKenna, A., Hanna, M., Banks, E., Sivachenko, A., Cibulskis, K., Kernytsky, A.,…DePristo, M. A. (2010). The genome analysis toolkit: A MapReduce framework for analyzing next-generation DNA sequencing data. Genome Research, 20(9), 1297–1303. doi:10.1101/gr.107524.110

Mcmahon, B. J., Teeling, E. C., & Höglund, J. (2014). How and why should we implement genomics into conservation? Evolutionary Applications, 7(9), 999–1007. doi:10.1111/eva.12193

Mohandesan, E., Speller, C. F., Peters, J., Uerpmann, H., Uerpmann, M., Cupere, B. E. A. D. E.,…Burger, P. A. (2017). Combined hybridization capture and shotgun sequencing for ancient DNA analysis of extinct wild and domestic dromedary camel. Molecular Ecology Resources, 17, 300–313. doi:10.1111/1755-0998.12551

Morin, D. J., Kelly, M. J., & Waits, L. P.(2016). Monitoring coyote population dynamics with fecal DNA and spatial capture–recapture. The Journal of Wildlife Management, 80(5), 824–836. doi:10.1002/jwmg.21080

Morin, P. A., Wallis, J., Moore, J. J., Chakraborty, R., & Woodruff, D. S. (1993). Non-invasive sampling and DNA amplification for paternity exclusion, community structure, and phylogeography in wild chimpanzees. Primates, 34(3), 347–356. doi:10.1007/BF02382630

Nichols, R. V., Konigsson, H., Danell, K., & Spong, H. (2012). Browsed twig environmental DNA: diagnostic PCR to identify ungulate species. Molecular Ecology Resources, 12, 983–989. doi:10.1111/j.1755-0998.2012.03172.x

Nielsen, E. E., Morgan, J. A. T., Maher, S. L., Edson, J., Gauthier, M., Pepperell, J.,…Ovenden, J. R. (2017). Extracting DNA from ‘jaws’: high yield and quality from archived tiger shark (Galeocerdo cuvier)skeletal material. Molecular Ecology Resources, 17(3), 431–442. doi:10.1111/1755-0998.12580

Oka, T., & Takenaka, O. (2001). Wild gibbons’ parentage tested by non-invasive DNA sampling and PCR-amplified polymorphic microsatellites. Primates, 42(1), 67–73. doi:10.1007/BF02640690

Orkin, J. D., Yang, Y., Yang, C., Yu, D. W., & Jiang, X. (2016). Cost-effective scat-detection dogs: unleashing a powerful new tool for international mammalian conservation biology. Scientific Reports, 6(1), 34758–34758. doi:10.1038/srep34758

Perry, G. H., Louis, E. E., Ratan, A., Bedoya-Reina, O. C., Burhans, R. C., Lei, R.,…Miller, W. (2013). Aye-aye population genomic analyses highlight an important center of endemism in northern Madagascar. Proceedings of the National Academy of Sciences of the United States of America, 110(15), 5823–8. doi:10.1073/pnas.1211990110

Perry, G. H., Marioni, J. C., Melsted, P., & Gilad, Y. (2010). Genomic-scale capture and sequencing of endogenous DNA from feces. Molecular Ecology, 19(24), 5332–44. doi:10.1111/j.1365-294X.2010.04888.x

Perry, G. H., Melsted, P., Marioni, J. C., Wang, Y., Bainer, R., Pickrell, J. K.,…Gilad, Y. (2012). Comparative RNA sequencing reveals substantial genetic variation in endangered primates. Genome Research, 22(4), 602–10. doi:10.1101/gr.130468.111

Perry, G. H., Reeves, D., Melsted, P., Ratan, A., Miller, W., Michelini, K.,…Gilad, Y. (2012). A genome sequence resource for the aye-aye (Daubentonia madagascariensis), a nocturnal lemur from Madagascar. Genome Biology and Evolution, 4(2), 126–35. doi:10.1093/gbe/evr132

Purcell, S., Neale, B., Todd-Brown, K., Thomas, L., Ferreira, M. A. R., Bender, D.,…Sham, P. C. (2007). REPORT PLINK: A Tool Set for Whole-Genome Association and Population-Based Linkage Analyses. The American Journal of Human Genetics, 81(September), 559–575. doi:10.1086/519795

Quéméré, E., Crouau-Roy, B., Rabarivola, C., Louis, E. E., & Chikhi, L. (2010). Landscape genetics of an endangered lemur (Propithecus tattersalli)within its entire fragmented range. Molecular Ecology, 19(8), 1606–21. doi:10.1111/j.1365- 294X.2010.04581.x

Rakoto-Joseph, O., Garde, F., David, M., Adelard, L., & Randriamanantany, Z. A. (2009). Development of climatic zones and passive solar design in Madagascar. Energy Conversion and Management, 50(4), 1004–1010. doi:10.1016/j.enconman.2008.12.011

Ralainasolo, F. B., Ratsimbazafy, J. H., & Stevens, N. J. (2006). Behavior and diet of the Critically Endangered Eulemur cinereiceps in Manombo forest, southeast Madagascar. Madagascar Conservation and Development, 3(1).

Randimbiharinirina, D. R., Raharivololona, B. M., Hawkins, M. T. R., Frasier, C. L., Culligan, R., Sefczek, T.,…Louis, E. E. (2017). Behavior and Ecology of male aye-ayes (Daubentonia madagascariensis)in the Kianjavato Classified Forest, southeastern Madagascar. Folia Primatologica, Manuscript.

Randimbiharinirina, D., Sefczek, T., Raharivololona, B. M., Andriamalala, Y., Ratsimbazafy, Jonah, & Louis, E. E. (2017). Aye-Aye Daubentonia madagascariensis (Gmelin, 1788)Madagascar (2016). In C. Schwitzer, R. A. Mittermeier, Rylands, Anthony B., F. chiozza, E. Williamson, E. Macfie,…A. Cotton (Eds.), Primates in Peril The world’s 25 most endangered primates 2016–2018.

Ravaloharimanitra, M., Ratolojanahary, T., Rafalimandimby, J., Rajaonson, A., Rakotonirina, L., Rasolofoharivelo, T.,…King, T. (2011). Gathering Local Knowledge in Madagascar Results in a Major Increase in the Known Range and Number of Sites for Critically Endangered Greater Bamboo Lemurs (Prolemur simus). International Journal of Primatology, 32(3), 776–792. doi:10.1007/s10764-011-9500-4

Robinson, J. T., Thorvaldsdottir, H., Winckler, W., Guttman, M., Lander, E. S., Getz, G., & Mesirov, J. P. (2012). Integrative Genomics Viewer. Nature Biotechnology, 29(1), 24–26. doi:10.1038/nbt.1754.Integrative

Sastre, N., Francino, O., Sanchez, A., & Ramirez, O. (2009). Sex identification of wolf (Canis lupus)using non-invasive samples, (July 2015. doi:10.1007/s10592-008-9565-6

Schmieder, R., & Edwards, R. (2011). Quality control and preprocessing of metagenomic datasets. Bioinformatics (Oxford, England), 27(6), 863–864. doi:10.1093/bioinformatics/btr026

Schubert, G., Stoneking, C. J., Arandjelovic, M., Boesch, C., Eckhardt, N., Hohmann, G.,…Vigilant, L. (2011). Male-mediated gene flow in patrilocal primates. PloS One, 6(7), e21514–e21514. doi:10.1371/journal.pone.0021514

Schwitzer, C., Mittermeier, R. a., Davies, N., Johnson, S. E., Ratsimbazafy, J. H., Razafindramanana, J.,…Nash, S. (2013). Lemurs of Madagascar: a strategy for their conservation 2013–2016 (No. 9781934151624).

Shafer, A., Wolf, J., Alves, P., Bergström, L., .MB., Brannstrom, I.,… Zielinski, P. (2015). Genomics and the challenging translation into conservation practice. Trends in Ecology & Evolution2, 30(2), 78–87. doi:10.1016/j.tree.2014.11.009

Sikes, R. S., & Gannon, W. L. (2011). Guidelines of the American Society of Mammalogists for the use of wild mammals in research. Journal of Mammalogy, 92(1), 235–253. doi:10.1644/10-MAMM-F-355.1

Smiley, T., Spelman, L., Lukasik-, M., Mukherjee, J., Ph, D. V. M. D., Kaufman, G.,…Cranfield, M. (2010). Noninvasive Saliva Collection Techniques for Free-Ranging Mountain Gorillas and Captive Eastern Gorillas E. Journal of Zoo and Wildlife Medicine, 41(2), 201–209.

Snyder-mackler, N., Majoros, W. H., Yuan, M. L., Shaver, A. O., Gordon, J. B., Kopp, G. H.,…Alberts, S. C. (2016). Efficient Genome-Wide Sequencing and Noninvasively Collected Samples. Genetics, 203(June), 699–714. doi:10.1534/genetics.116.187492

Sterling, E. J. (1994a). Aye-aye: Specialists on Structurally Defended Resources. Folia Primatologica, 62, 142–154.

Sterling, E. J. (1994b). Taxonomy and Distribution of Daubentonia madagascariensis: A Historical Perspective. Folia Primatologica, 62, 8–13.

Sterling, E. J., & McCreless, E. E. (2007). Adaptations in the Aye-aye: A Review. In Lemurs (pp. 159–184).

Sundqvist, A.-K., Ellegren, H., & Vila, C. (2008). Wolf or dog? Genetic identification of predators from saliva collected around bite wounds on prey. Conservation Ge, 9, 1275–1279. doi:10.1007/s10592-007-9454-4

Tan, C. L. (1999). Group composition, home range size, and diet of three sympatric bamboo lemur species (genus Hapalemur) in Ranomafana National Park, Madagascar. International Journal of Primatology, 20(4), 547–566.

Valiere, N., & Taberlet, P. (2000). Urine collected in the field as a source of DNA for species and individual identification, 2150–2153.

Wheat, R. E., Allen, J. M., Miller, S. D. L., Wilmers, C. C., & Levi, T. (2016). Environmental DNA from Residual Saliva for Efficient Noninvasive Genetic Monitoring of Brown Bears (Ursus arctos). PloS One, 1–18. doi:10.5281/zenodo.58748

Wright, P. C., Johnson, S. E., Irwin, M. T., Jacobs, R., Schlichting, P., Lehman, S.,…Tan, C. (2008). The Crisis of the Critically Endangered Greater Bamboo Lemur (Prolemur simus), 2008(23), 5–17.

